# PubNavigator as an undergraduate teaching tool

**DOI:** 10.1101/2023.09.18.558293

**Authors:** Rosario A. Marroquín-Flores, Maverick A. Campbell, Stephanie Grimes, Lisa B. Limeri

## Abstract

The ability to use data and evidence as the basis for scientific reasoning is a critical skill that undergraduate students must learn to successfully transition into science careers. There is a need for pedagogical tools that promote the development of science process skills and reduce barriers to implementation in the classroom. PubNavigator (www.pubnavigator.com) is a science communication platform that shares the findings of primary literature using accessible language and features biographical information about the authors. This research assesses the utility of PubNavigator as an undergraduate teaching resource. We analyzed responses to an assignment prompting discussion of PubNavigator articles in an asynchronous online discussion forum in an introductory biology course. We conducted thematic analysis guided by the evidence-based reasoning framework to identify evidence of science reasoning and interactive enagement. We found evidence that the PubNavigator assignment elicited engagement in scientific reasoning. Students who demonstrated higher levels of engagement with the assignment also demonstrated higher levels of science reasoning in their discussion of the articles. Our findings support PubNavigator as an undergraduate teaching tool that can be used to promote science reasoning and engagement in introductory biology. We provide recommendations for implementation to instructors interested in integrating PubNavigator activities into their courses.

## INTRODUCTION

Vision and Change (2011) calls for undergraduate biology education to move beyond content and to help students develop science process skills (American Association for the Advancement of Science, 2011). Science process skills, such as reading primary literature and using higher-level reasoning, are necessary abilities for those working in science careers. Students must develop strong science process skills to successfully transition into science careers and many instructors recognize the value of teaching science process skills in the classroom (Coil et al., 2010). However, there are several obstacles that prevent instructors from explicitly teaching these skills, with time constraints being chief among them (Brownell & Tanner, 2012; Coil et al., 2010; Petersen et al., 2020; Shadle et al., 2017). There is an urgent need to develop pedagogical tools that promote the development of science process skills, and which can easily integrate into existing curricula.

Science reasoning, the ability to use evidence and data as the basis for scientific arguments, is a science process skill that helps students create new knowledge and evaluate scientific claims (Brown et al., 2010; Coil et al., 2010). Prior work suggests that group work and peer interaction may be effective tools for promoting scientific reasoning (Chi & Wylie, 2014; Halmo et al., 2022). Chi and Wylie (2014) developed a framework of cognitive engagement in learning, in which interacting with others’ ideas is the highest level that best promotes student learning (ICAP framework). Halmo and colleagues (2022) analyzed the conversations of students engaged in a problem solving activity using the Evidence-Based Reasoning Framework and identified an association between metacognitive regulation skills and higher-level science reasoning (Brown et al., 2010; Halmo et al., 2022). These findings suggest that group work can promote scientific reasoning in undergraduate STEM students by prompting them to respond to each other and justify their arguments using evidence (Halmo et al. 2022). Reading primary literature has also been shown to promote science reasoning and the development of other science process skills in students (Abdullah et al., 2015; Hoskins et al., 2007; Kozeracki et al., 2006; Krontiris-Litowitz, 2013). The wide adoption of the CREATE (<u>C</u>onsider, <u>R</u>ead, <u>E</u>lucidate hypotheses, <u>A</u>nalyze and interpret data, and <u>T</u>hink of the next <u>E</u>xperiment) model, in which students read several primary research articles from a single laboratory, suggests that instructors recognize the value of integrating primary literature in the classroom (Hoskins et al., 2007). While effective, many of the existing approaches require instructors to take time to alter the course and to work with students, who often struggle with the complex language and concepts found in primary literature (Janick-Buckner, 1997; Van Lacum et al., 2012). There is a need for alternative approaches that reduce barriers to implementation in the classroom.

There are some extant tools that have been used to make primary literature more accessible to novice readers (Goudsouzian & Hsu, 2023). For example, Figure Facts is a Microsoft Word template that contains a blank table that students can complete as they read the primary article, designed to help students to focus on the data figures and tables (Round & Campbell, 2013). Figure Facts can be used to improve data interpretation skills and reduce the stress associated with reading primary literature (Round & Campbell, 2013). Annotated Primary Literature is an additional tool that adds context to previously published articles (Kararo & McCartney, 2019; McCartney et al., 2018). The articles are housed on an online platform and have clickable “layers” added to improve scaffolding to promote student engagement and understanding (Kararo & McCartney, 2019; McCartney et al., 2018). An alternative approach that has frequently been used in secondary classrooms is to integrate Adapted Primary Literature, which retains the structure of a primary research article, but is modified by teachers to meet the needs of a novice reader and can be adjusted to fit within the classroom context (Falk et al., 2008; Koomen et al., 2016; Yarden et al., 2001, 2015). Researchers have taken several approaches to integrating Adapted Primary Literature into the classroom and findings suggest that they can facilitate scientific discourse, help students to understand the nature of science, and have been recommended to support the development of science reasoning skills (Ariely et al., 2019; Department of Agriculture and Forestry Service, 2013; Falk et al., 2008; Koomen et al., 2016). However, Adapted Primary Literature relies on teachers to self-author the adapted text and the extensive amount of work required by the instructor limits its utility as an undergraduate educational tool that can be widely adopted.

This research assesses the utility of a novel Adapted Primary Literature tool as an undergraduate teaching resource. PubNavigator (www.pubnavigator.com) is a science communication platform that hosts articles that feature the findings of primary literature using accessible language and features biographical profiles of the researchers who authored the article. We assess the capacity for PubNavigator articles to provide an accessible introduction to primary literature and foster interactive engagement with primary literature. PubNavigator articles do not retain the structure of the primary research article, but instead, summarize a primary paper in several sections. The “Research at a Glance” section provides an overview of the research, similar to an abstract. The “Highlights” section contains a detailed description of one primary finding or method. The “What my Science Looks Like” section includes an image or diagram that shows part of the research process, often including the researchers themselves. Each article is written at or below a 12^th^ grade reading level and contains a “Decoding the Language” section with definitions for science jargon used within the body of the article, often paired with examples. Each article also includes a “Learn More” section, which provides links to government databases, websites, and/or YouTube videos that readers can follow to learn more about the topic of the article. Importantly, all PubNavigator articles also include biographical information about the focal researcher, with the goal of making the scientist behind the science more accessible and relatable. The biographical section includes pictures of the author and information about the authors’ research field, study organism, career goals, and hobbies. More recent articles also include information about how the focal researcher became a scientist. The PubNavigator biographies were designed based on prior research which has demonstrated that classroom activities that help undergraduate students relate to scientists lead to increased interest in science, reduced stereotypes about scientists, improved academic outcomes, and can promote science identity formation (Aranda et al., 2021; Brandt et al., 2020; Ovid et al., 2023; Schinske et al., 2016; Yonas et al., 2020).

We developed an online PubNavigator assignment that can be easily integrated into existing undergraduate biology curricula with minimal time investment from instructors. Here, we explore the capacity for PubNavigator to support the development of science process skills through science reasoning and interactive engagement with peers about research. We implemented this lesson in an introductory biology course and analyzed student assignment responses to assess the extent to which PubNavigator reading assignments motivate students to engage with the activity and foster students’ practice of science process skills.

## METHODS

We conducted a qualitative study to assess the effectiveness of assignments using PubNavigator articles in an introductory biology course at a public R2: Doctoral-granting institution. The introductory biology course primary enrolls freshman-level students and is designed for biology majors. Undergraduate students enrolled in the course were assigned three PubNavigator articles to read and discuss as part of a laboratory assignment. Their responses were qualitatively analyzed to evaluate scientific reasoning and interactive engagement. At the time of the study, all students were attending courses remotely due to covid-related university closures. Recorded lectures were available to students at all times. The class met synchronously online once per week for the lecture section to work through examples and discuss questions regarding the material. The laboratory sections also met synchronously once per week for brief lectures on the laboratory material and to discuss the weekly at-home experiments. All procedures were approved by the Illinois State University Institutional Review Board (#IRB-2020-181).

### Assignment

The assigned PubNavigator articles highlighted research conducted by graduate and undergraduate students in the School of Biological Sciences at Illinois State University. The articles were selected to align with topics in the course curriculum. The first article highlighted research on the Beneficial Acclimation Hypothesis, where researchers sought to understand the effect of temperature on the prevalence and intensity of parasitic infections in the common Eastern Bumble Bee (*Bombus impatiens*) (Tobin et al., 2019). The second article highlighted research on inbreeding depression, where researchers examined mating success and offspring production in inbred and outbred populations of the decorated cricket (*Gryllodes sigillatus*) (Sakaluk et al., 2019). The final article highlighted research on temperature-dependent sex determination, where researchers examined the effects of heat waves and fluctuating incubation temperatures on male and female development in the red-eared slider turtle (*Trachemys scripta*) (Carter, 2018). The students were asked to read the articles and discuss them in an online, asynchronous forum as part of a laboratory assignment. On the discussion board, students were placed into groups of 5-7 students and groups were changed for each assignment. Students were encouraged to discuss anything that they found interesting about the article but were given several prompt questions to help initiate discussion (Figure 1).

**Figure 1.**
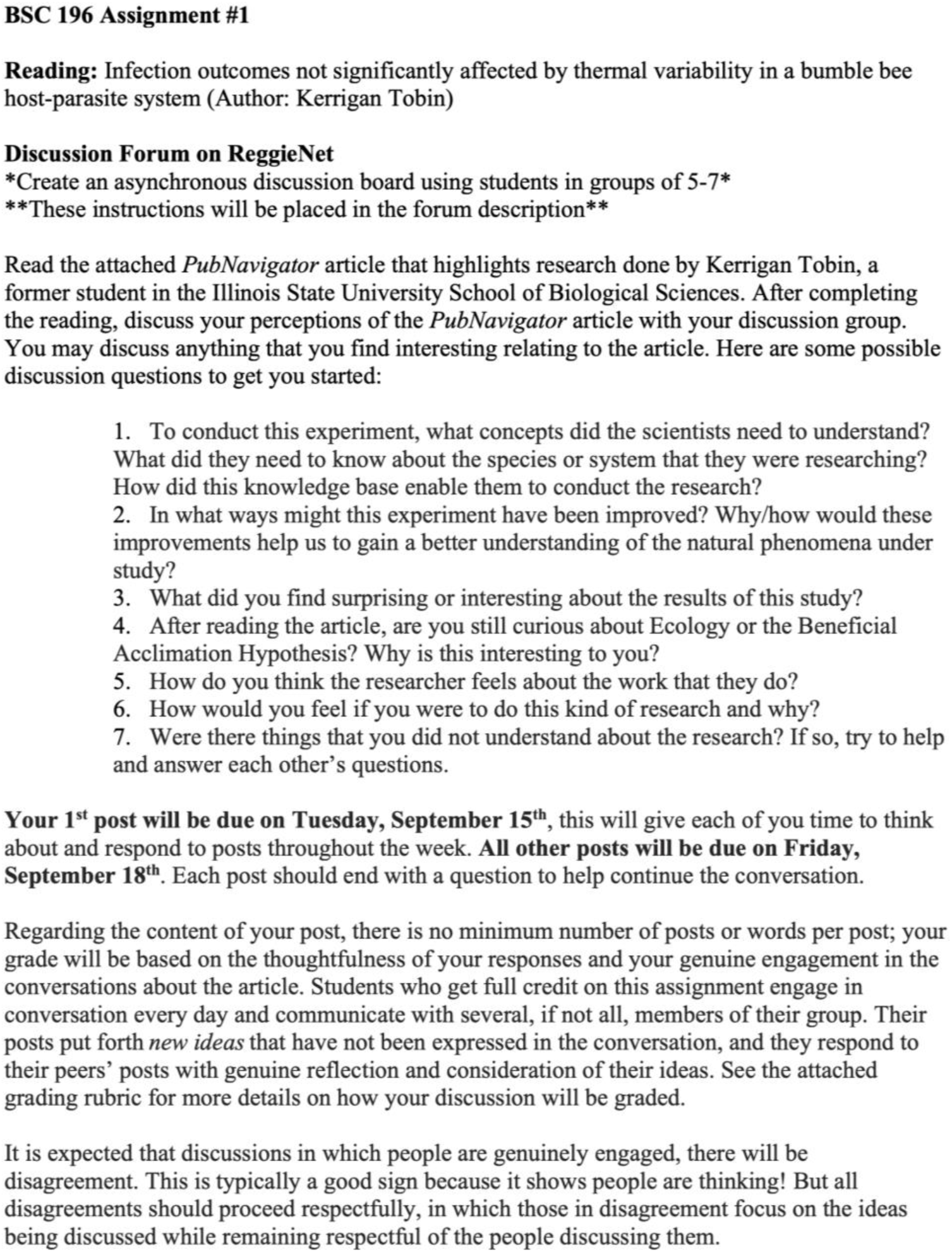
Example of the PubNavigator assignment including prompt questions, due dates, and expectations.

The PubNavigator assignment was graded based on the timeliness and thoughtfulness of the initial post, engagement with other students, and contribution to the discussion (Table 1). The asynchronous discussion board posts were collected for analysis at the end of the semester for students who consented to participate in the study.

**Figure 1.** Example of the PubNavigator assignment including prompt questions, due dates, and expectations.

| **Grading Rubric (15 points)** |  |  |  |
| --- | --- | --- | --- |
| **Initial Post**<br>(2 points) | • Posts well-developed, fully addresses all aspects of the task.<br>• Post is made on time.<br>(2 points) | • Posts are adequate, may include superficial thought and preparation; do not address all aspects of the task.<br>(1 point) | • No post is included.<br>(0 points) |
| **Engagement**<br>(5 points) | • Posts to the discussion with clear, concise comments daily.<br>• Engages with most, if not all, other contributing group members.<br>• Ends each post with a question to prompt further discussion.<br>(5 points) | • Posts to the discussion with clear, concise comments.<br>• Engages with a small number of contributing members.<br>(3 points) | • Posts are long, unorganized, or include rude content.<br>(1 point) |
| **Contribution**<br>(8 points) | • Genuinely responds to posts and poses new questions to engage further discussion.<br>• Uses references to the literature, readings, or personal experiences to support comments.<br>• Contributes to and broadens the discussion.<br>(8 points) | • Posts and responses are adequate, may include superficial thought and preparation.<br>• Uses references to personal experiences, but no references to other research or other readings.<br>(5 points) | • Posts are long, unorganized, or include rude content.<br>(0 points) |

### Participants

Consent was obtained on a pre-semester survey. Two-hundred and fifty-seven students were enrolled in the course and all students completed the assignment as a regular part of the course. Only students who consented to participate and completed all three assignments were included in data analysis (n = 23). Demographic data is not available for the participants in this study.

### Characterizing Science Reasoning and Reading Engagement

In our qualitative analysis of student discussion board posts, we took a thematic approach, the process of identifying and analyzing patterns within the data (Braun & Clarke, 2006). We conducted data analysis in two phases. In the first phase, we used a deductive approach to characterize science reasoning and in the second phase we used an inductive approach to characterize reading engagement. Each phase involved an iterative process in which three members of the research team (R.A.M.F., M.A.C., S.G.) first coded text (i.e., read a section of text and identified themes present in the text, labeled as “codes” identifying the themes) individually, then met to discuss the codes applied to each section of text to consensus. Group discussions among the analysts would occasionally result in changes to the codebook or coding approach, in which case all previously coded data was revisited with the new approach/codebook. All qualitative data were analyzed using Delve Coding Software (https://delvetool.com/).

### Phase 1: Deductive analysis of Scientific Reasoning

In the first phase of our analysis, we used the Evidence-Based Reasoning Framework to deductively evaluate students’ ability to use evidence as the basis for scientific reasoning (Braun & Clarke, 2006). The approach allowed us to describe the relationship between basing arguments in evidence and making correct statements about scientific concepts. The analysis approach was iteratively revised over time, which we describe in three steps.

The Evidence-Based Reasoning Framework was used as the basis for defining the components of a reasoning unit (Brown et al., 2010). Similar to other work, a complete reasoning unit was defined as a statement that included a premise and a claim, linked by some form of support (Halmo et al., 2022). Support was defined as the use of data, evidence, or a rule (Brown et al., 2010; Halmo et al., 2022).

#### Step 1

In our initial coding, we allowed statements that included an implied rule to be considered a complete reasoning unit. An implied rule was not explicitly stated but could be deduced based on the context of the premise and claim. For example, after reading the article on temperature-dependent sex determination, one student wrote, “Similar to my response to [classmate], I think with warmer weather, both scenarios are possible. I think the population would ultimately decrease and mating would be limited…”. In this example, the claim is that climate change will lead to a reduced population of turtles and that it will be more difficult for turtles to find mates. The implied rule for this statement is that higher temperatures lead to more female turtles, which was the subject of the assigned reading.

Metacognitive statements have been linked to higher-level reasoning in group work (Halmo et al., 2022). To explore the relationship between quality of reasoning and metacognition, we further classified reasoning units in terms of correctness, level of peer interaction, and completeness (Halmo et al., 2022). A statement was coded as “correct” if it accurately conveyed a scientific idea described in the reading or made accurate comparisons to other scientific information outside of the reading. A statement was coded as “mixed” when it included a combination of correct and incorrect ideas about a scientific concept (Halmo et al., 2022). Each reasoning unit was coded as correct or mixed, even if the reasoning unit was incomplete. This approach allowed us to make comparisons across all reasoning units. A statement was coded as “transactive” when a student was directly responding to another student within the forum. Transactive statements were identified when students used statements, such as, “I agree with [student]” or “Like [student] said,”. All other statements were coded as “non-transactive.” A statement was coded as “complete” when it included at least one form of support for a claim, and “incomplete” when it made a claim without any forms of support.

#### Step 2

In the second phase of coding, the definition of a reasoning unit was refined to better fit the context of the assignment being assessed in this study. In the original definition, a reasoning unit had to contain a premise to be complete. However, we found that students were frequently referencing the same reading or discussion prompt prior to making a claim, and thus the premise was implied. Requiring the premise to be explicitly stated was limiting our ability to capture students’ enacted reasoning. Therefore, we revised the definition of a complete reasoning unit to require only a claim and a form of support (Figure 2). We also removed statements that included an implied rule from the complete reasoning units. Including implied rules involved a leap in logic wherein the research team had to identify potential sources of support that students were trying to use as a basis for their arguments, which required the researchers to make the arguments for the students. Previously coded data were re-analyzed based on these changes.

**Figure 2.**
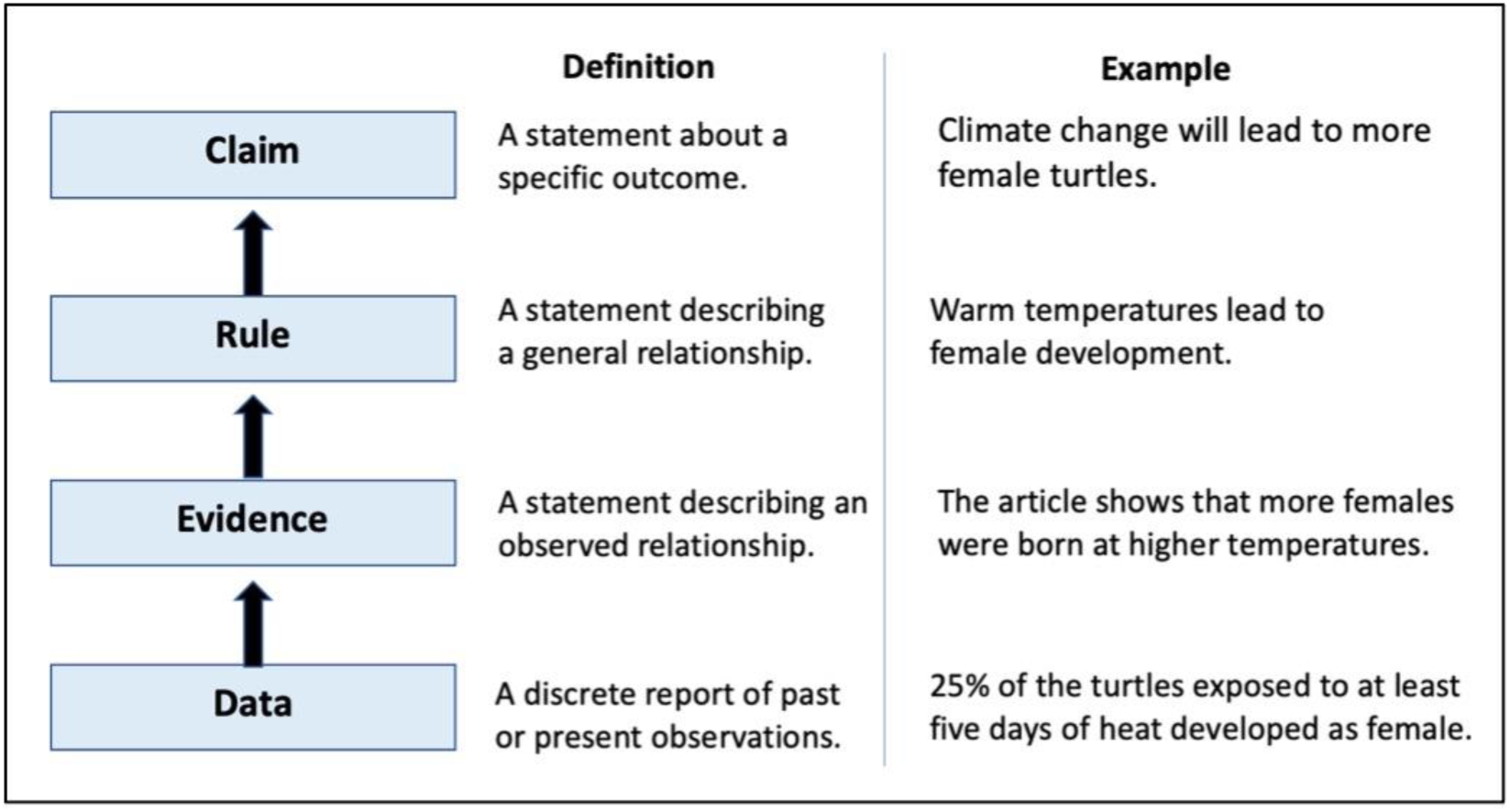
Example of a complete reasoning unit (Brown et al., 2010).

#### Step 3

After we coded all reasoning units, we noticed that the reasoning units varied in the amount of support that was included in defense of each claim. Some students were using more support while others were using less, or none. Therefore, we further characterized each reasoning unit to identify reasoning units with high, intermediate, and low support. Reasoning units with high support were defined as a unit including two or more forms of support to justify an argument (i.e., the student used data and evidence to support a statement). Reasoning units with intermediate support were defined as a reasoning unit including only one form of support to justify an argument (i.e., the student only used evidence or only used data to support a statement). Reasoning units with low support were defined as units which contained no forms of support to justify an argument. Since low support reasoning units lacked support, they were also considered incomplete.

After further characterizing the reasoning units, we identified a trend; students that used more support also tended to make more correct statements. This observation led us to question why students were making mixed statements. We posited two potential explanations: 1) students were making mixed statements because the PubNavigator article did not properly convey information from the primary article, or 2) students did not engage with the reading, and thus, were not prepared to accurately discuss the scientific concepts described within the articles. To address our first potential explanation, we began the process of combining and splitting codes to better understand why students were making mixed statements (Saldaña, 2013). Mixed statements were broken into subunits that included “misunderstanding of a biological concept”, “misunderstanding of the article”, “misunderstanding of outside information”, and “general misconceptions.” Our approach allowed us to capture the full breadth of why students were making mixed statements. To address our second potential explanation, we proceeded to the second phase of the qualitative analysis, characterizing student engagement with the assignment.

### Phase 2: Inductive analysis of Engagement

In the second phase of our analysis, we used an inductive approach to identify additional themes that emerged from the data (Braun & Clarke, 2006). The approach allowed us to describe the relationship between student engagement and science reasoning and to describe student perceptions of the assignment. Similar to phase 1, phase 2 involved iterative revisions to the codebook and approach, which we describe in three steps below.

#### Step 1

We used an inductive approach to identify evidence of engagement. Unlike our approach to characterizing scientific reasoning, we did not have a pre-existing codebook for analyzing student engagement. Coding for engagement was a highly iterative process where codes were created, combined, and deleted to better fit the research question. Members of the research team individually coded each instance that a theme appeared in the data, then met to come to consensus on each code. This process was repeated for each new theme that emerged from the data.

Our initial approach to measuring engagement was to identify evidence of “reading enjoyment” and “connecting with the research and the scientist” because we anticipated that learning about authentic research conducted by graduate students within the department would spark an interest in the field and help students connect with the researcher. However, we identified patterns in the data based on student understanding and effort and began to question whether students were making an effort to understand the reading. To address this question, we added an additional set of codes: “importance of the article”, “critique of the article”, and “connecting the content within the article to other coursework”.

#### Step 2

After we had coded all the student responses, we revisited all coded excerpts to critically re-evaluate whether all codes represented a single idea and were distinguishable from each other. Each coder independently read all coded excerpts for each code and made notes. The research team then met to discuss any issues identified, with the goal of breaking down codes that contained multiple ideas into grandchild codes and combining codes that could not be distinguished. We also removed some codes that did not pertain to our initial research questions (e.g., “off-topic conversations”). Our code “connecting the content within the article to other coursework” was expanded to include other types of connections (termed “making connections”). Students would often relate the findings of the articles to their own personal experiences, such as a family trip or weather in their hometowns. Students also often compared the information in the article to other biological concepts. For example, many students reading the paper about temperature-dependent sex determination in turtles made comparisons to sex-determination in other species. We also added an interest code that was further split into different types of interest: “learned something new”, “desire to participate”, and “general interest”. Our approach allowed us to identify students who 1) were connecting information from the article to their existing knowledge base, 2) demonstrated an interest in the reading, and 3) the ways in which students were demonstrating their interest.

#### Step 3

In our final review of the codebook, we opted to change the names of some codes to more precisely describe the meaning of the code and make the distinctions between each code more evident. For example, the code named “making connections”, where students connect information from the article to other knowledge and experiences, was renamed as “relating to outside knowledge”. The new name is more descriptive of the meaning of the code and more distinguishable from a separate code, “connecting to the scientist”, where students described an emotional connection to the focal researcher. The finalized codebook included parent, child, and grandchild codes describing dimensions of scientific reasoning and student engagement (Table 2).

**Table 2.**
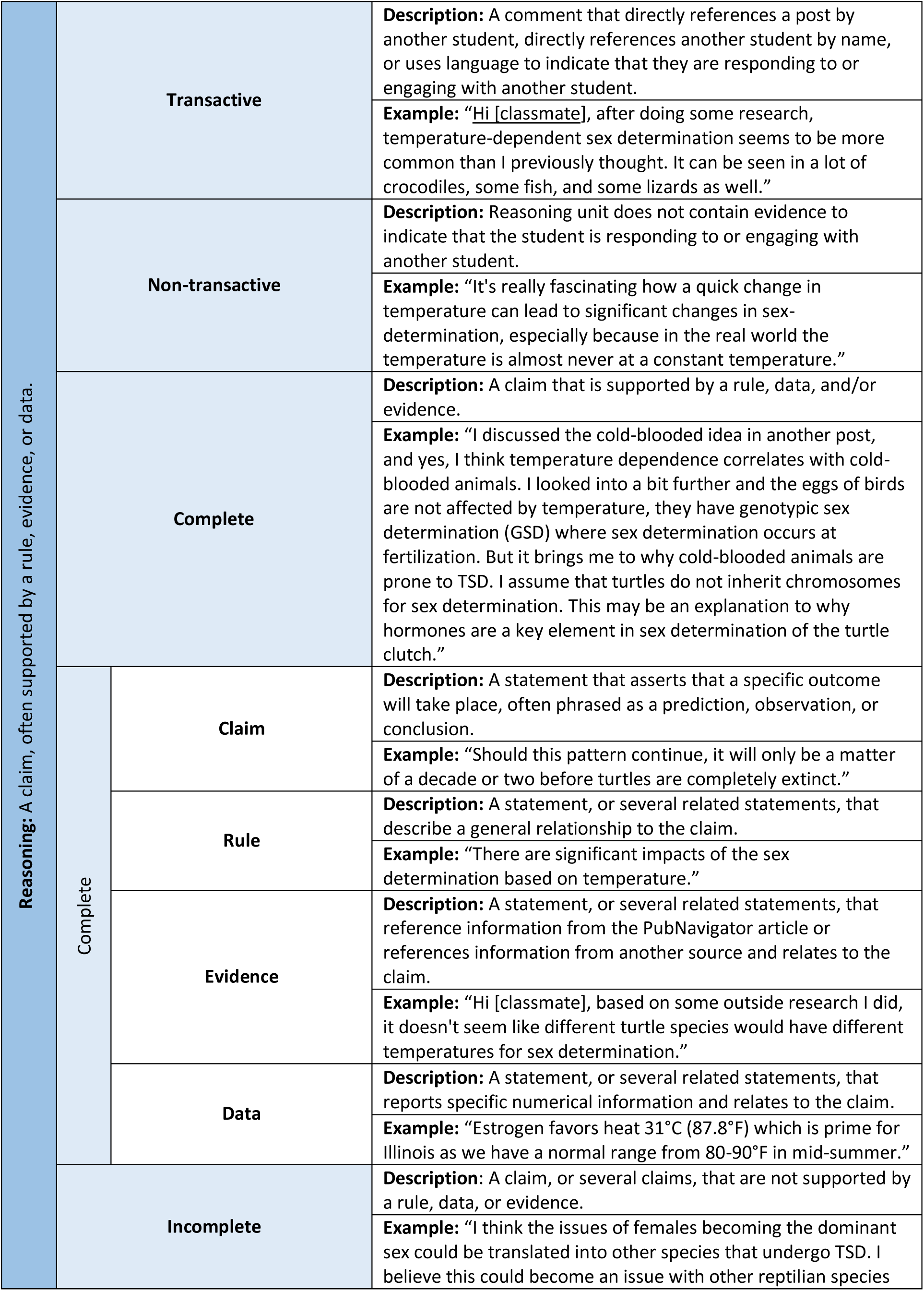
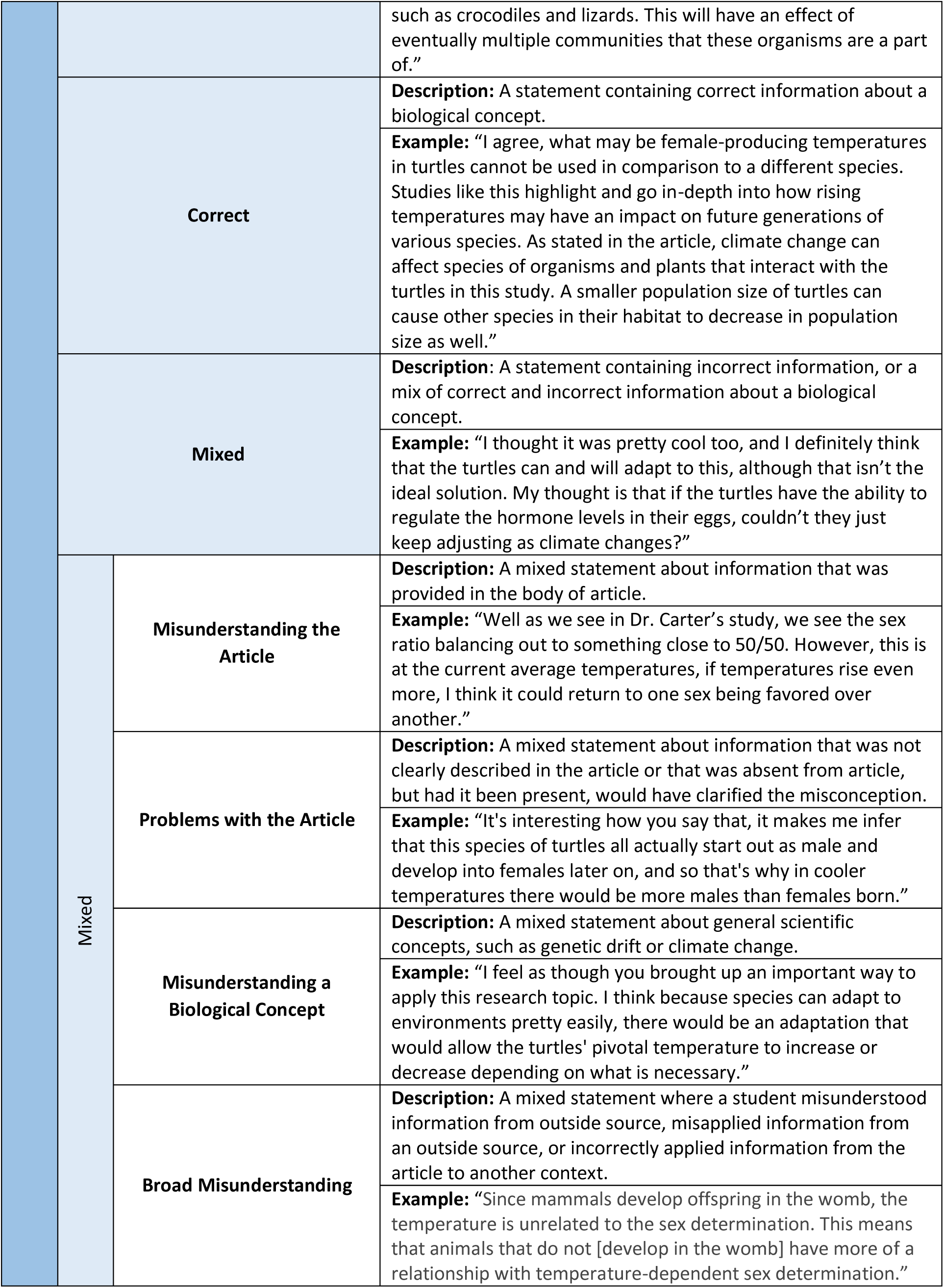
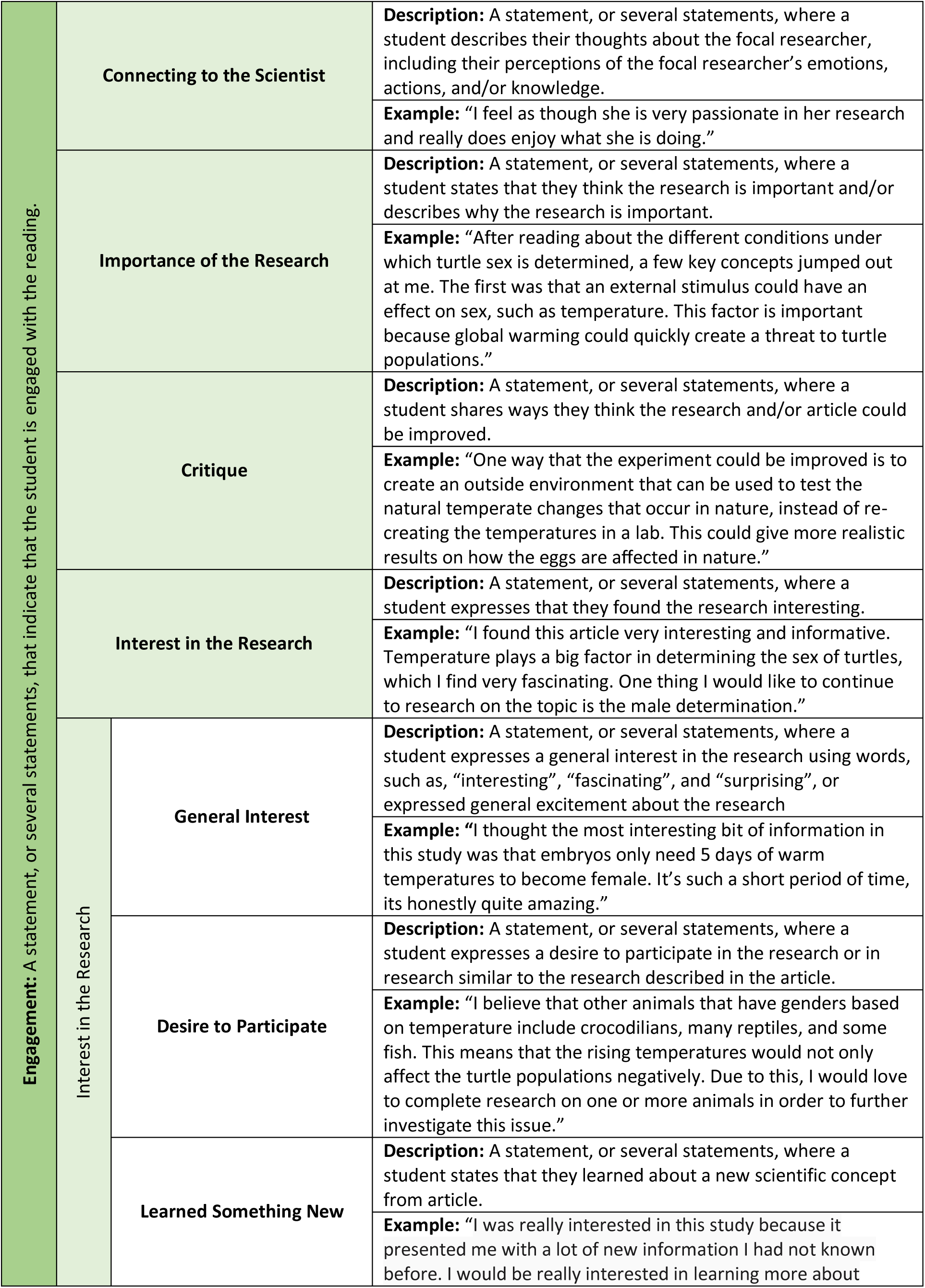
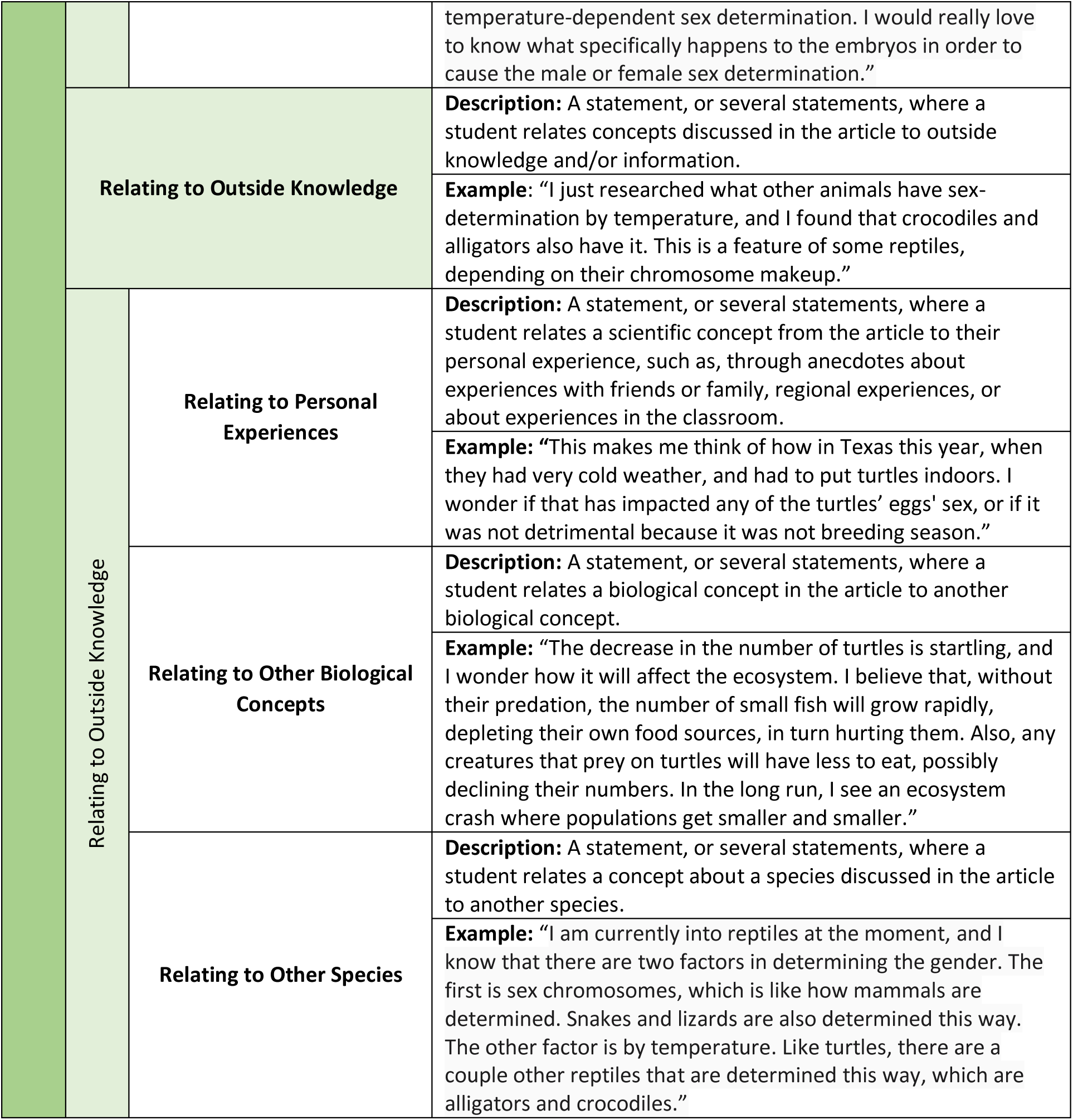
Full codebook for science reasoning and engagement. The student quotes are lightly edited for grammar, spelling, and typographical errors to improve readability.

### Characterizing the Relationship between Science Reasoning and Reading Engagement

Prior research suggests that higher-level reasoning is associated with complete, correct, and transactive statements (Halmo et al., 2022). Based on our findings from the qualitative data analysis, we decided to further explore the relationships between our science reasoning codes and reading engagement. To accomplish this, we quantified the number of overall reasoning units and overall engagement units in each assignment. We then used the Kruskal-Wallis test to determine if there was a difference in the number of reasoning and engagement units between assignments. Next, we grouped students based on their use of support for each assignment. Students who had at least one reasoning unit with high support within an assignment were placed into the “high support” category. Students who had at least one reasoning unit with intermediate support within an assignment were placed in the ‘intermediate support” category. Students who did not have any complete reasoning units within an assignment were placed into the “low support” category. We then used the Kruskal-Wallis test to determine if there was a difference in the number of reasoning and engagement units based on the amount of support that students used to justify their claims.

We also quantified the number of complete, transactive, and correct reasoning subunits. To control for variation in the number of reasoning units made by students, we calculated for each participant the proportion of all their statements which were categorized as complete, transactive, and correct. We then used the Kruskal-Wallis test to determine if there was a difference in the proportion of correct, complete, or transactive reasoning units by assignment or by support category. We did not have sufficient data to quantitatively determine the relationship between different types of engagement (engagement subunits) and assignment or based on the amount of support that students used. Quantitative analyses were conducted using R Studio (Version 2023.06.1+524) (Core Team, 2022).The Wilcoxon rank sum test was used for all *post hoc* tests.

In assignments 2 and 3, one student had substantially more reasoning and engagement units than the other students. Additionally, three students remained in the same support category across all three assignments. The analysis described above was conducted both with these students included and with these students removed from the data. Removing these students did not change the results of the analyses. Therefore, the findings presented below retain all participating students.

## RESULTS

### Science Reasoning

All participating students demonstrated some form of science reasoning (Tables 2-3). However, the number of reasoning units differed by assignment and by student support category. There was a significant difference in the number of reasoning units by assignment (χ^2^ = 8.7469, df = 2, p = 0.013), where students made significantly more reasoning units in the final assignment compared to the first assignment (p = 0.0074; Figure 3). There was no difference in the number of reasoning units between the second and third assignments (p = 0.295).

**Figure 3.**
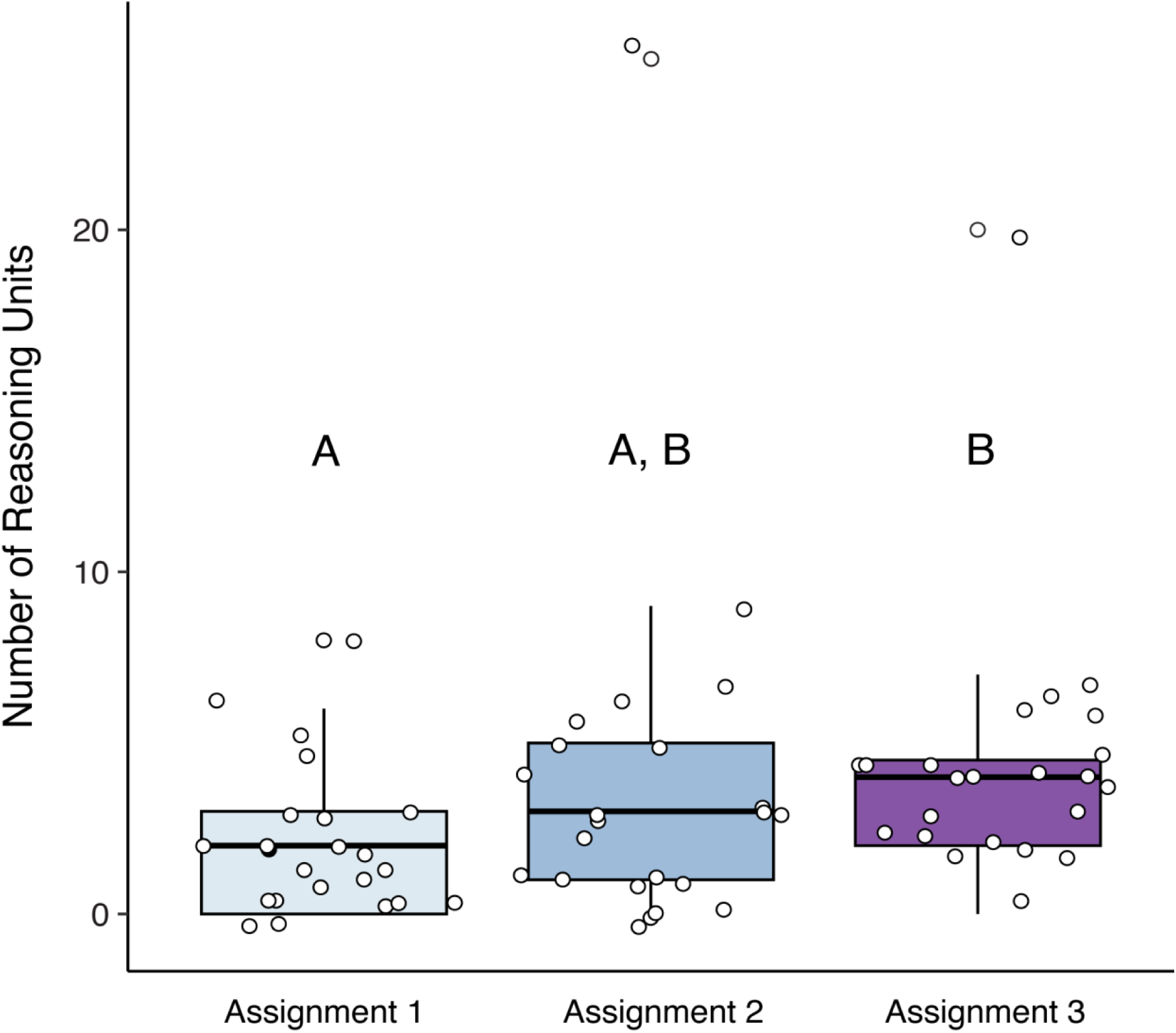
The total number of reasoning units that students made in each assignment (Assignment 1 = Beneficial Acclimation Hypothesis in bumblebees, Assignment 2 = Inbreeding depression in crickets, Assignment 3 = temperature-dependent sex determination in turtles). Different letters denote significant differences.

**Table 3.** Child and grandchild codes for science reasoning with example exerpts of student responses. Formatting is added to the student responses for complete and incomplete reasoning units to indicate the codes assigned. The student responses are lightly edited for grammar, spelling, and typographical errors to improve readability.

| Reasoning | Examples of student statements |
| --- | --- |
| **Complete reasoning unit with high support**<br>22 students had at least one complete reasoning unit with high support. | **<u>“Inbreeding has lots of negative effects.”</u>** To agree with you again, I would have loved to know why these negative effects occur. *In my opinion, which is purely thought, not scientifically backed up, these negative effects may occur due to the health-harming genes in a family having a greater opportunity than before (with mates not in their family) to come to light. For example, if a family carries a disease, you have the chance to not get it once mating with a partner who does not have said disease. However, once you inbreed, you make it much more likely to see this disease.”*<br><u>Data, Evidence, Rule, Claim</u> |
| **Complete reasoning unit with intermediate support**<br>22 students had at least one complete reasoning unit with intermediate support. | *“I would personally guess that it was either some specific trait of the parasite or the temperature difference wasn't enough to cause a drastic bodily response in the bees. <u>When converted to Fahrenheit, the bees only endured an overall temperature difference of 14 degrees.</u> To me, that number does not seem like it would make too much of a difference in the bees' body to resisting the parasite.”*<br><u>Data, Evidence, Rule, Claim</u> |
| **Incomplete reasoning unit (low support)**<br>23 students had at least one incomplete reasoning unit | *“As far as I know, some other reptile species such as crocodiles and other turtle species have temperature-dependent sex determination. I also was very fascinated by the idea of TSD, as it is something that I haven't learned much about throughout my lifetime. Typically, sex is determined by the sex chromosomes given by each parent, but sex chromosomes seem to not be the deciding factor when a species has TSD.”*<br><u>Data, Evidence, Rule, Claim</u> |
| **Transactive reasoning unit**<br>21 students had at least one transactive reasoning unit. | Example 1: “To try to answer your question, I believe that the animals that are inbreeding in nature will not do it intentionally, due to most populations being spread out and having lots of diversity. Rather, I think that inbreeding occurs when a population is in decline and there are only a few individuals left.”<br>Example 2: “I think that, like [classmate] said, it would depend on which animal we would be using in another experiment because not all animals determine sex like turtles do”. |
| **Non-transactive reasoning unit**<br>22 students had at least one non-transactive reasoning unit. | Example 1: “For this experiment, it was important to know the conditions in which crickets would mate. Most animals and insects have specific mating rituals or behaviors. To produce the best results, you would need to be able to replicate the wild mating conditions.”<br>Example 2: “Typically, sex is determined by the sex chromosomes given by each parent, but sex chromosomes seem to not be the deciding factor when a species has TSD.” |
| **Correct reasoning unit**<br>21 students had at least one correct reasoning unit. | Example 1: “I'm sure the inbred crickets are still genetically similar to the outbred crickets, however that population probably has less genetic variation due to inbreeding.”<br>Example 2: “I think that the research chose a small range of temperature because climate change is gradual. Climate change does not happen rapidly and all at |
|  | once, so seeing the effect of more temperatures may not be as realistic. I think that I bigger change would have different results though.” |
| **Mixed reasoning unit**<br>21 students had at least one mixed reasoning unit. | Example 1: “I like that you researched how they respond to temperature! This changes things completely, even if the inbred crickets were low in numbers <u>increasing reproduction rates by chirping more could produce more crickets.</u> Despite the difficulties with reproduction this idea might encourage reproduction within the outbred and inbred crickets.”<br>Example 2: “His research had <u>proven that the BAH was not accurate</u> when looking at bees fighting against parasitic infections in different climates.” |

There was also a significant difference in the number of reasoning units based on student support category (χ^2^ = 36.433, df = 2, p << 0.001, Figure 4). Students in the low support category had significantly fewer reasoning units than those in the intermediate support (p << 0.001) and high support (p << 0.001) categories. Similarly, students in the intermediate support category had significantly fewer reasoning units than those in the high support category (p = 0.006).

**Figure 4.**
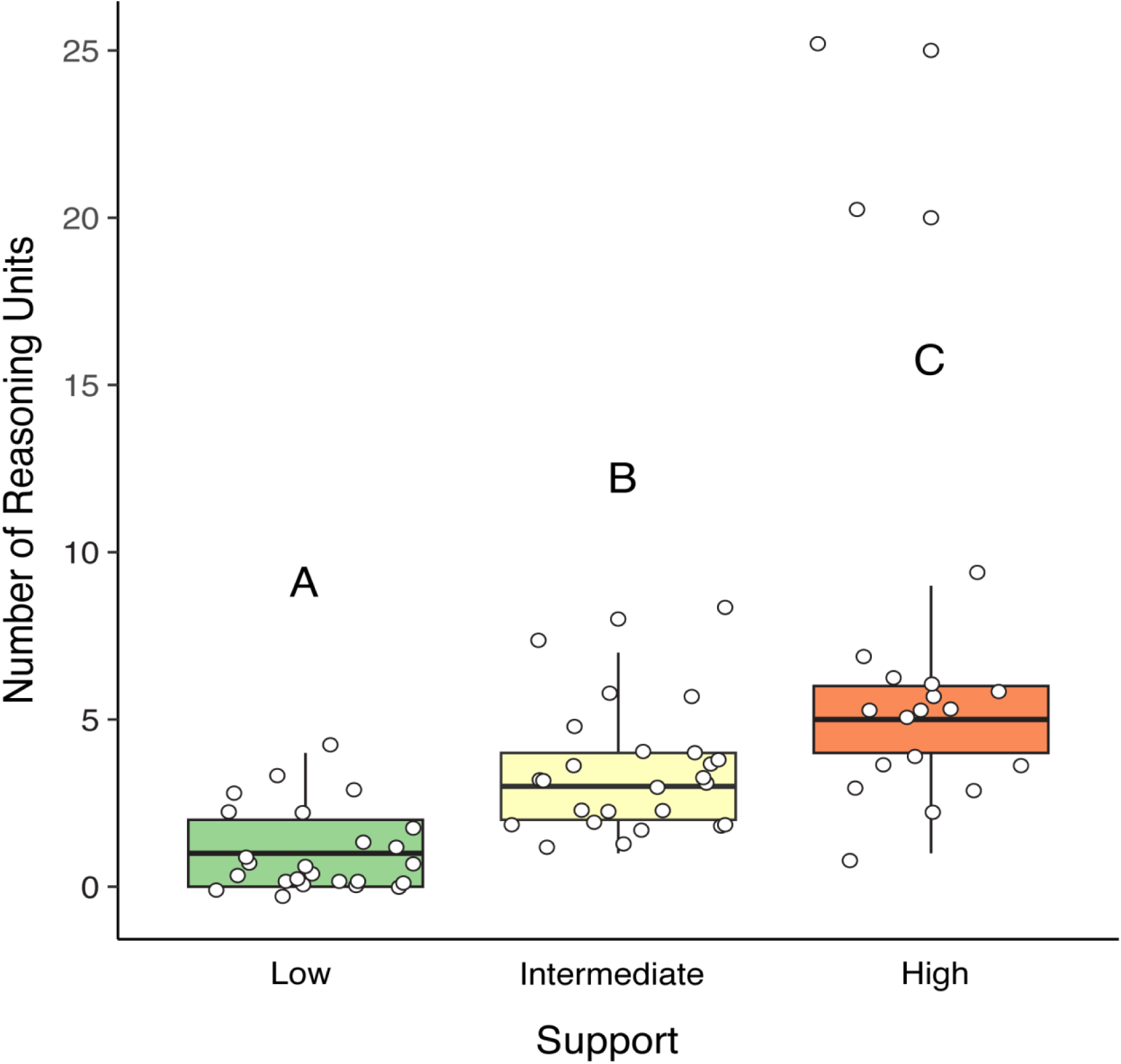
The total number of reasoning units made by students in each support category across the three assignments. Different letters denote significant differences.

We next examined the relationship between correct or transactive statements and the strength of support that students used. To control for variation in the number of reasoning units made by students, we calculated the proportion of reasoning units made by each student that were correct and transactive. We found a significant relationship between the proportion of correct reasoning units by student support category (χ^2^ = 19.292, df = 2, p << 0.001; Figure 5), where students in the low support category made significantly, proportionately fewer correct reasoning units than those in the intermediate (p < 0.001), and high (p < 0.001) support categories. There was no significant relationship between the proportion of transactive reasoning units and support category (χ^2^ = 5.0631, df = 2, p = 0.080).

**Figure 5.**
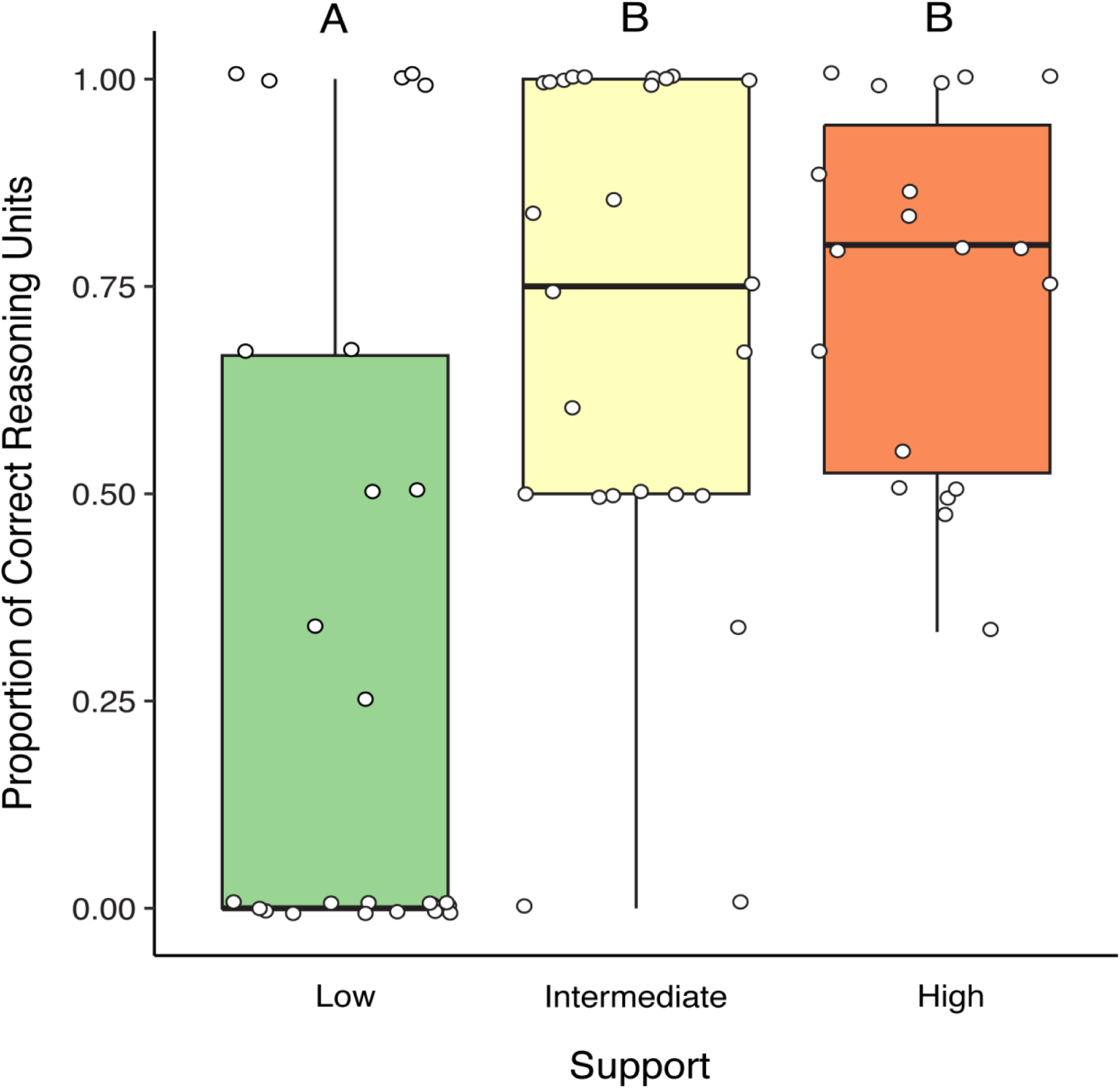
The proportion of correct reasoning units made by students in each support category across the three assignments. Different letters denote significant differences.

### Why are students making mixed statements?

In our analysis of the mixed reasoning units, we found that incorrect ideas rarely seemed to come from the article itself, but rather were related to misconceptions about a biological concept and more general misconceptions related to the life-history traits of the animals under study and the impacts of species decline. Across these misconceptions, we found several similarities in the way that students were thinking about biology. Cognitive construals refer to the informal and intuitive ways that students commonly think about the world, and which apply to a student’s understanding of biological concepts (Coley & Tanner, 2015, 2012). The misconceptions identified in the mixed reasoning units showed evidence of common cognitive construals.

One cognitive construal is teleological thinking, a type of causal reasoning where students assume that there is a goal, purpose, or function to the biological concept under consideration (Coley & Tanner, 2015, 2012). In our results, we observed that students often used teleological thinking in their discussions of evolution, adaptation, and mating. Six students demonstrated teleological thinking in their discussion of the PubNavigator articles. For example, one student wrote:

> I find your theory to be interesting and myself believe that the release of estrogen into the embryos could also be a result of evolution. I agree that the point of this evolutionary event to occur is to avoid only male turtles being born which could potentially cause extinction.

In this quote, the student exemplifies teleological thinking by describing an evolutionary event as directional, having the goal or purpose of preventing extinction of the species.

Another cognitive construal is anthropocentric thinking, a tendency to reason about unfamiliar biological concepts by making analogies to humans (Coley & Tanner, 2015, 2012). In the context of this study, anthropocentric thinking often manifested in the personification of animals and the natural world. Seven students demonstrated anthropocentric thinking in their discussion of the PubNavigator articles. For example, one student wrote:

> If there were more regular males recently introduced, I think it would take a while for the inbred females to realize that these males would produce healthier offspring--considering if they have not been exposed to these regular males, they may not be able to recognize them right away.

In this quote, the student is assigning human characteristics to a non-human organism (Coley & Tanner, 2015). Rather than responding to attractive physical traits that may be related to adaptative qualities, the student suggests that female crickets are actively evaluating prospective mates based on their perceived capacity to create healthy offspring.

Another type of cognitive construal is essentialist thinking, implicit assumptions that there is some form of essential property common to all members of a given category (Coley & Tanner, 2015). A consequence of essentialist thinking is the perception that all category members are uniform and predictable, which often leads students to discount variability in nature (Coley & Tanner, 2015). Eight students demonstrated essentialist thinking in their discussion of the PubNavigator articles. For example, one student wrote:

> Normally, I would assume that due to the inbreeding nature of these crickets, the species would be led to extinction because the number of offspring (with short lifespans and damaged health) exponentially decreases with every generation; however, these crickets seem to contradict our expectations. Regardless, I am going to stick with my assumption and believe that the population would eventually be led to extinction.

In this quote, the student assigns innate, negative, characteristics to all inbred crickets. The student suggests that inbred critckets have no innate potential and that they will all experience unavoidable deleterious effects as a result of inbreeding. However, one finding in the article was that female crickets from the inbred lines preferentially selected inbred males over outbred males as mates. The authors stated in the article that, in light of these findings, “…we cannot simply conclude that ‘inbreeding is bad’…”. Despite this information, the student insists that the innate and unavoidable deleterious effects of inbreeding will lead the population to extinction.

### Reading Engagement

All participating students demonstrated some form of engagement with the activity and there was no difference in the number of engagement units by assignment (χ^2^ = 0.47868, df = 2, p = 0.787). In our analysis of the relationship between reasoning and engagement, we found a significant difference in the number of engagement units by support category (χ^2^ = 19.448, df = 2, p << 0.001; Figure 6). Students in the low support category had significantly fewer engagement units than students in the intermediate (p = 0.002) and high (p < 0.001) support categories. There was no difference in the number of engagement units between students in the intermediate and high support categories (p = 0.068).

**Figure 6.**
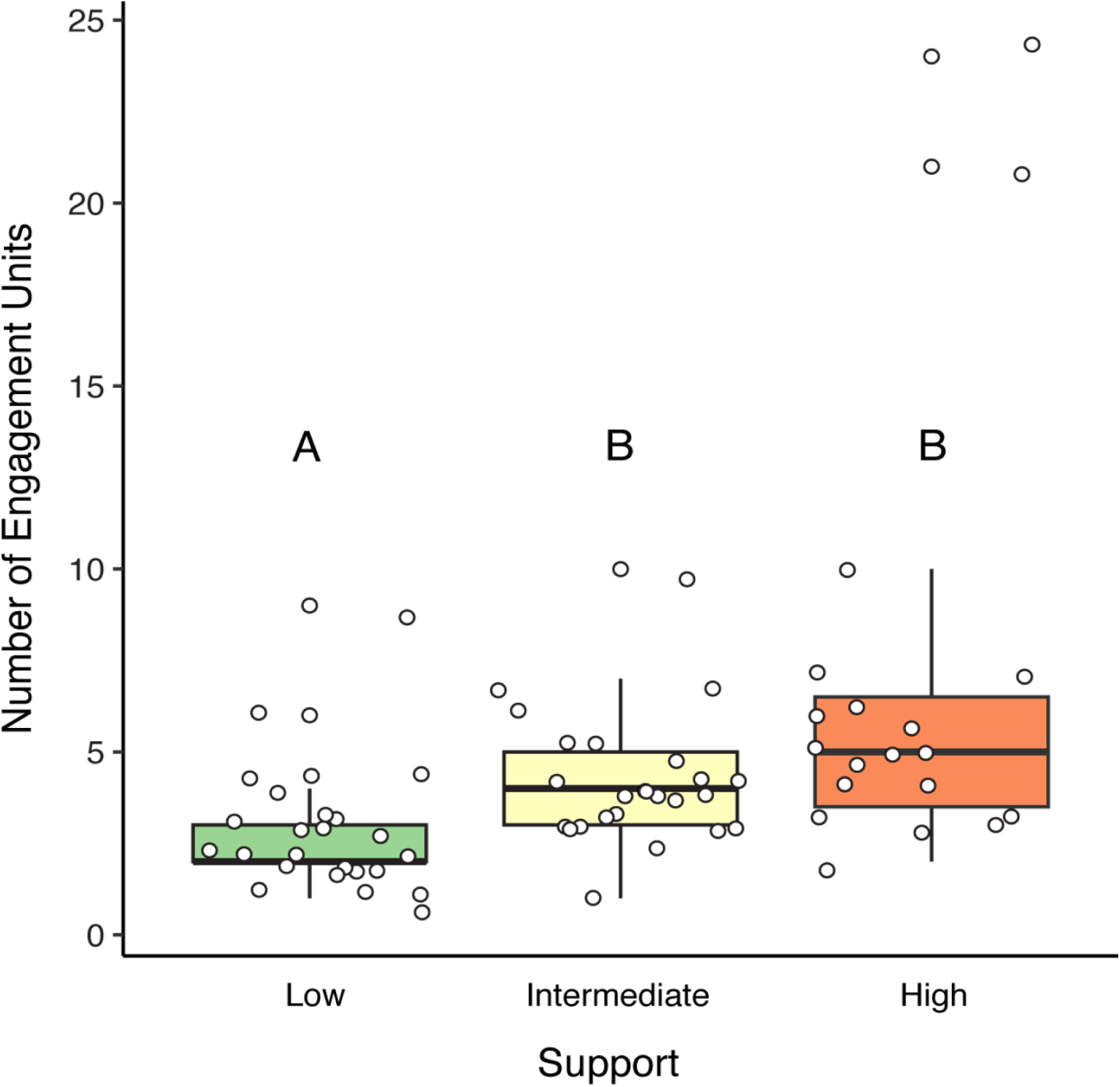
The total number of engagement units made by students in each support category across the three assignments. Different letters denote significant differences.

In our qualitative analysis, we identified several ways in which students were engaging with the content (Table 2). Students primarily engaged by relating the content to outside knowledge, demonstrating interest in the reading, discussing its importance, offering critique of the experiment, and by connecting with the scientist highlighted in the article.

#### Relating to outside knowledge

Almost all students related the content from the assignments to some form of outside knowledge. We identified three ways in which students connected the material to their experiences: 1) connecting the information from the article to other species (n = 19), 2) relating the information to other biological content knowledge (n = 20), and 3) relating the information to their personal experiences (n = 13). Students who were connecting the information to other species were often doing so to better understand the findings of the article and the system under study. For example, one student wrote:

> My understanding of genetics in turtles is very limited, but if it is anything like human genetics, another factor that could potentially be at play is the SRY gene. The SRY gene is a part of the Y chromosome that, to put it simply, makes an organism into a male. Perhaps a higher temperature could affect whether this SRY gene in males is initiated or inhibited.

In this quote, the student uses their knowledge about about sex determination in mammals to better understand temperature-dependent sex determination in turtles. The student posits that a Y-linked gene responsible for the initiation of testis development in mammals may also be related to male development in turtles. Rather than coming from the Y-chromosome, as is does in mammals, the student suggests that temperature may be responsible for regulating the expression of SRY in male turtles.

Students relating the information to other biological concepts were often referencing mechanisms of evolution or discussing trophic interactions to better understand the outcomes of the research. One such student wrote:

> I believe there will be complications not only within the population, but also with other populations that require the survival of the population affected by inbreeding. For example, if the crickets from these experiments were unable to reproduce healthy offspring, that could end up affecting the survival of predators that feed on those crickets. Needless to say, inbreeding can lead to a chain reaction of multiple populations being negatively affected.

In this quote, the student discusses the potential negative impacts of inbreeding despression. The student describes the ecological connectedness between different organisms. Crickets can serve as prey items for predators and low prey availabaility can have negative impacts on predator populations, and thus, can have cascading effects on the ecosystem.

Other students attempted to predict the long-term effects of the findings of the research. For example, one student wrote:

> Since climate change is occurring at such a slow rate, I do believe that there is a possibility that the eggs would eventually adapt to the changes in temperature in time. Evolution might help produce more offspring that can possibly survive which can help future generations. The offspring may vary in their appearance, physiology and behavior, and some of those variants are better able to survive than others. Therefore, over succeeding generations, the appearance of organisms will demonstrate new adaptations. However, evolution takes place within generations, therefore, adaptations aren’t guaranteed.

In this quote, the student tries to predict how turtles with temperature-dependent sex determination will respond to climate-related changes in temperature over time. The student suggests that the population may adapt to changing temperatures due to the development of beneficial traits that improve survival and reproduction over time.

Students who related the information from the articles to their personal experiences primarily referenced experiences in school or experiences specific to the region. For example, one student wrote:

> In one of the lectures in this class, it was hypothesized that females prefer their mates to be healthy and more likely to produce healthier offspring (which is suggested by the brighter the color of a male, the healthier they are). However, in this experiment the females tended to prefer mating with males that will be more likely to produce less healthy offspring. The ideas the researchers made for this phenomenon do make sense, I just find it interesting that this experiment differed from the general hypothesis made in the biology lecture.

In the quote, the student compares one of the findings from the paper to information that they learned about sexual selection in a biology course. The student comments on the surprising differences between the findings of the article on inbreeding depression in crickets and what they learned in class.

#### Interest

All students expressed some form of interest while completing the assignments. We identified three ways in which students demonstrated their interest: 1) expressing general interest in the reading (n = 23), 2) discussing something new that they learned from the reading (n = 19), and 3) describing a desire to participate in the research presented in the article or in similar research (n = 13). Students expressing general interest in the research were often expressing enjoyment with the reading or stating that they found the reading “interesting.” In both the article on the beneficial acclimation hypothesis in bumblebees and the article on temperature-dependent sex determination in turtles, students were often surprised to learn about the range of effects that climate change can have on living organisms and how they interact with one another. For example, one student wrote:

> I was super surprised at the results of Carter’s experiment because I would have never thought that temperature directly affects the gender of an embryo and whether it becomes a male or female. I had a little knowledge that various environmental factors would affect the development of embryos in eggs but never imagined climate could control something such as the sex.

In this quote, the student was surprised to learn that temperature can influence gonadal differentiation in animals and developed a more complete understanding of the potential impacts of climate change on temperature-sensitive species.

In the article about inbreeding depression in crickets, many students were learning about inbreeding depression for the first time, while others were surprised to learn more about the life history traits of crickets. One student wrote, “After reading this article, I am interested in learning more about inbreeding depression because it is something that I have not learned about before, and I think it’s interesting that traits worsen as those same traits are bred together.” In this quote, the student describes learning about inbreeding depression for the first time and describes an interest in learning more about the topic.

Students expressing a desire to participate in the research often described an interest in doing research in a lab setting and expressed more interest in working with turtles and climate change than working with the invertebrates. For example, one student wrote:

> I would be very interested in performing this study as it shows how a rise in temperature can affect different animal populations. This is due to how I’m personally interested in how the earth and the animals are being affected by the rise in temperatures.

In this quote, the student describes an existing interest in learning about the effects of climate change. The article aligned with the student’s interests and made them interested in conducting similar research.

#### Importance

Almost all the students discussed the importance of the research presented in at least one of the articles. Fewer students discussed the importance of the research conveyed in the article on inbreeding depression in crickets (n = 8) compared to the articles on the beneficial acclimation hypothesis in bumblebees (n = 14) and temperature-dependent sex determination in turtles (n = 12). However, students who did discuss the importance of inbreeding depression focused on how the research could be used to inform conservation efforts. Those discussing the other two articles primarily addressed the importance of understanding the effects of climate change. For example, one student wrote, “Research like this is important because it helps us to see why pollinators are disappearing. Our work will help to focus conservation efforts. Human-driven climate change has many effects on earth, but full impacts are not well understood.” In this example, the student describes how the findings of the study can be used to respond to climate change and promote conservation.

#### Critique

Nearly all the students engaged with the reading by offering a critique of the research contained in the article. In the article on the beneficial acclimation hypothesis in bees, the students most often critiqued the number of variables in the study, which consisted of two strains of bumblebee parasite, a range of incubation temperatures, and several colonies of bees. For example, one student wrote:

> To improve this, I would have used the same exact parasite instead of two different ones and used bees all from the same colony, and tested them in two different temperatures. I do not quite understand why two different parasites were used for this experiment, it should be assumed that every parasite, infection and sickness will not affect the host the same or last the same duration of time, as well as begin at exactly the same time. What I would do might create a more accurate outcome.

In this quote, the student describes how they would improve upon the experiment if they were to conduct it themselves. The student points out some potential variability that may have been introduced into the experiment by using multiple colonies and strains of parasite. The student argues that reducing these points of variability would improve the quality of the data. Students also critiqued the range of temperatures used, arguing that more extreme temperatures would more closely mimic the effects of climate change.

In the article on inbreeding in crickets, student critiques focused on ways to expand the study to better understand the findings. One such student wrote:

> I think they should have further emphasized the behaviors and processes of the crickets that were not inbreeding. I found it helpful that they included this on some of their graphs, but not all. In order to understand how they differ, we need to understand the underlying behaviors.

In this quote, the student suggests adding another source of data collection. In the article, the authors described watching crickets mate in a mating chamber illuminated by red light. The authors then measured hatching success, development time, and the number of offspring produced by inbred and outbred mated pairs (Sakaluk et al., 2019). Here, the student suggests adding some of the observational data from the mating chamber to see if there were other differences between inbred and outbred crickets.

In the article on temperature-dependent sex determination in turtles, students most frequently suggested expanding the study to include several generations of turtles, or to run a field experiment to more closely mimic the conditions that a turtle would experience in nature.

#### Connecting with the scientist

Fourteen of the participating students made comments relating to the researcher highlighted in the article. Students who connected with the highlighted scientists most frequently described how passionate they thought the author was and how they imagined the author perceived their work. For example, one student wrote:

> I think the timing of this research spotlight discussion couldn’t be more perfect as I am currently writing speeches in my communications class analyzing global warming and climate change. One thing I found interesting then and after reading this research article is that many of the effects of climate change are unknown. For that reason, I think that Dr. Amanda Wilson Carter would be excited to contribute research to such a serious topic, as would I.

In this quote, the student connects the content of the article to their own experiences and their own learning. The student suggests that learning about climate change is very important and projects their excitement about the topic onto the focal researcher. The perceived excitement of the focal researcher comes from the student’s own excitement about the research. Another student wrote:

> I think that Tobin was confident in her work. The way that she described her processes and findings was very thorough and detailed. Her research had proven that the BAH was not accurate when looking at bees fighting against parasitic infections in different climates. If I were to do this kind of research, I think I would also feel confident in my findings.

Similar to the prior example, the student projects their own feelings about the research onto the scientist. If the student were to conduct this research, they would be confident in their approach to data collection and analysis, therefore, the focal researcher must also be confident.

## DISCUSSION

Introductory-level STEM courses represent an entry point into science majors, but frequently serve as “weed out” courses (Meaders et al., 2020; Mervis, 2011). Student experiences in introductory courses can influence their decision to remain in STEM majors (Christopher Strenta et al., 1994; Meaders et al., 2020; Seymour & Hewitt, 1997; Watkins & Mazur, 2013). Many instructors continue to use teacher-centered pedagogy in introductory courses, and thus, students are rarely introduced to the process of science in introductory courses (Coil et al., 2010; Ebert-May et al., 2011; Petersen et al., 2020; Shadle et al., 2017). We have developed a low-stakes assignment that can be easily integrated into introductory-level biology courses and that encourages students to engage in science reasoning. The goal of this study was to assess the efficacy of PubNavigator assignments in promoting student engagement in science reasoning skills and interactive engagement with classmates. We found that students engaged in science reasoning and interacted with their peers in the process of completing PubNavigator assignments.

Halmo and colleagues (2022) defined higher-level reasoning as transactive, complete, and correct, arguing that the exchange of ideas between students can lead to higher quality reasoning. Similarly, Chi and Wylie (2014) state that activities that challenge students to interactively generate new ideas achieve the greatest level of learning and that the mutual exchange of ideas must occur at regular intervals to truly be considered interactive. In our PubNavigator assignment, we asked students to discuss their reading in an online discussion and students were graded based on engagement and genuine contribution to the discussion. The assignment instructions and assignment structure were designed to encourage students to interact with each other daily. Our approach was effective in motivating students to regularly contribute to the online discussion board. We found evidence of engagement among all our participating students and that students practiced science reasoning as they engaged in discussion.

The ability to use scientific evidence to support scientific claims sets the foundation for scientific thinking (Coil et al., 2010; Kuhn, 1993). All students demonstrated some form of scientific thinking across all three assignments, but the reasoning units differed based on the amount of support that students used to justify their claims. The most complex reasoning units were those that contained several forms of support to justify a given claim. In our analysis, we found a relationship between reading engagement and the strength of students’ arguments. Students who were more engaged with the assignment used more support to justify their claims. Students who used more support to justify their claims also made more correct statements about biological concepts. Surprisingly, we did not find a relationship between support and the transactive nature of student discussions. While students were encouraged to respond to one another daily, the nature of student interaction likely differs in an asynchronous discussion forum compared to an in-person activity. For example, rather than receiving immediate feedback in an in-person setting, students had to wait for their peers to respond on the platform. Without the ability to quickly build off each other’s arguments, students most frequently posted new ideas or topics, supported by evidence from external readings, personal experiences, or from the article itself.

### Identifying with the highlighted researcher

Research using Scientist Spotlights suggests that weekly assignments that encourage students to learn about counter-stereotypical scientists, and to reflect on their learning, lead to better academic outcomes and increase students’ ability to relate to the focal researcher (Schinske et al., 2016). In the PubNavigator assignment, we asked students to read a brief biography about the focal researcher and students were encouraged to reflect on the reading. We anticipated that students would better connect with focal researcher because each of the researchers highlighted in the PubNavigator articles were graduate students conducting research within the same department in which the course was situated. In the initial Scientist Spotlight assignments, nearly 80% of students were able to personally relate to an important scientist by the end of the semester, often due to shared interests with the scientist or personal qualities about the scientist (Schinske et al., 2016). In our implementation of the PubNavigator assignments, 61% of the students connected to the researcher highlighted in the article. However, none of the students explicitly mentioned the author’s status as a graduate student in their department or referred to the hobbies and interests described in the biography. Students primarily identified with the enthusiasm that the highlighted researcher displayed, often referring to researcher’s “passion” for the project.

There are several potential explanations for these differences. First, we did not explicitly measure scientist relatability. Instead, we looked for qualitative evidence to suggest that students were connecting with the highlighted researcher. Second, students spent more time learning about the highlighted researcher in the Scientist Spotlight assignments compared to PubNavigator assignments. In Scientist Spotlight assignments, students were asked to engage with content outside of the biography that introduced the personal history of the highlighted researcher, such as by viewing a TED Talk or listening to a podcast about the researcher (Schinske et al., 2016). Finally, the researchers highlighted in the PubNavigator articles did not hold, or did not explicitly disclose, minoritized or intersectional identities. In the assignment, we selected articles to cover content that fit within the course curriculum, without consideration of the identity of the focal researcher. Each of the PubNavigator articles highlighted at least one female researcher, but none of the highlighted researchers identified as a person of color, nor did they self-disclose unseen identities. While the highlighted researchers were graduate students within the department, the research took place at a primarily white institution, where students and faculty of color are critically underrepresented. It is possible that featuring counter-stereotypical examples of scientists, like those featured in Scientist Spotlights, may be more effective at improving the relatability of the highlighted researcher than featuring near-peers.

### Intuitive reasoning in student discussions

Students often enter introductory classrooms with fundamental misconceptions about biological concepts (Coley & Tanner, 2015, 2012; Stern et al., 2018). Many students reading the PubNavigator articles had mixed reasoning units and the inaccuracy of their statements primarily originated from misconceptions about a biological concept, rather than a misinterpretation of the reading or a problem with the article itself. Similar to other work, we identified several instances of both teleological and anthropocentric thinking in relation to the topics of evolution and natural selection in student discussions (Gregory, 2009; Pickett et al., 2022; Richard et al., 2017). Students often referred to evolution and natural selection as deterministic; traits evolved with direct goal of increasing the chances of survival and reproduction (i.e., teleological thinking). Students also often anthropomorphized the study organism or nature itself, describing individual animals as having goals (e.g., in mate selection) or how nature prevents species decline. While some authors have argued that anthropomorphism is a type of teleological thinking, we have presented them separately to distinguish between the types of misconceptions that emerged from our analysis (Gouvea & Simon, 2018). We also identified several instances of essentialist thinking, where students described an innate or unchanging property of living organisms (Coley & Tanner, 2015, 2012). Statements that contained essentialist thinking were particularly common in the discussions of the article on inbreeding depression in crickets. Students often disregarded individual variation and assumed that all inbred populations were absent of genetic diversity, that evolution was unachievable, and that all inbred populations would die over time. As these types of misconceptions are often associated with types of intuitive reasoning, it is unlikely that students would be able to resolve these misconceptions in an online discussion (Coley & Tanner, 2015, 2012; Pickett et al., 2022). Instead, instructors may choose to address these misconceptions in the classroom.

### Recommendations for implementation

The PubNavigator platform contains several articles that address concepts related to evolution and natural selection. Given the prevalence of common misconceptions regarding these topics in introductory classrooms, it may be beneficial to pair PubNavigator reading assignments that address evolution and natural selection with activities that explicitly address these misconceptions. One option could be to integrate refutation text into lecture as a think-pair-share or into a small group discussion held during lab. Refutation text refers to passages that include a commonly held misconception followed by an explicit refutation of the misconception with a scientific explanation for why it is incorrect (Tippett, 2010). Previous research suggests that students who read refutation text use less language associated with cognitive construals and show improved conceptual understanding of biological evolution (Asterhan & Resnick, 2020; Pickett et al., 2022).

### Limitations of the work

Our research represents the first step in evaluating PubNavigator as an undergraduate teaching tool. We found that PubNavigator assignments elicited scientific reasoning and engagement with research articles among students in an introductory biology course. While this initial evidence is promising, there are notable limitations to our current study that present opportunities for future research. We took a qualitative approach to measuring science reasoning and engagement with a small population of students. Due to the sample size and the absence of demographic data, we caution a broad application of these findings. The absence of demographic data limits our capacity to disaggregate student responses and better understand how students who hold minoritized or intersectional identities connect with the highlighted researchers. Future work should incorporate quantitative measures of scientist relatability, reading self-efficacy, and/or science identity to better understand student outcomes after engaging with PubNavigator assignments.

### Conclusion

PubNavigator is a science communication platform that communicates the results from primary literature in an accessible way. We have used articles from the PubNavigator platform to create a new teaching tool that encourages students to practice scientific reasoning and engage with their peers. We analyzed student responses to the assignment and found evidence that the assignment successfully elicits engagement in scientific reasoning among undergraduates in an introductory biology course. We also found that students who demonstrated a higher level of engagement with the articles also made more correct and more complete statements about the articles in their discussion. PubNavigator reading assignments can be selected to fit within the curriculum and can be easily be integrated into an online or in-person lab associated with the course.

## Acknowledgements

This research was made possible through a microgrant awarded to R.A.M.F. from the Sai Resident Collective. We would like to thank the Honors College Undergraduate Research Scholars Program supported by The <u>CH</u> Foundation and the Helen Jones Foundation, Inc. We would like to thank Dr. Rebekka Darner at the Illinois State Univeristy Center for Mathematics, Science, and Technology for her support and guidance during early data collection. We would like to thank Dr. Victoria Borowicz and Dr. Diane Byers at the Illinois State University School of Biological Sciences for supporting research within their course. We would also like to thank Dr. Rachael DiSciullo, Austin Calhoun, Dr. Daniel Goldberg, Elyse McCormick, Elliot Lusk, Jaclyn Everly, and Sydney Metternich, the Graduate Teaching Assistants responsible for grading the PubNavigator assignments.

## References

Abdullah, C., Parris, J., Lie, R., Guzdar, A., & Tour, E. (2015). Critical Analysis of Primary Literature in a Master’s-Level Class: Effects on Self-Efficacy and Science-Process Skills. CBE—Life Sciences Education, 14(3), ar34. 10.1187/cbe.14-10-0180

American Association for the Advancement of Science. (2011). Vision and change in undergraduate biology education: A view for the 21st century. Washington, DC. https://www.aaas.org/programs/inclusive-stemm-ecosystems-equity-diversity-iseed

Aranda, M. L., Diaz, M., Mena, L. G., Ortiz, J. I., Rivera-Nolan, C., Sanchez, D. C., Sanchez, M. J., Upchurch, A. M., Williams, C. S., Boorstin, S. N., Cardoso, L. M., Dominguez, M., Elias, S., Lopez, E. E., Ramirez, R. E., Romero, P. J., Tigress, F. N., Wilson, J. A., Winstead, R., … Tanner, K. D. (2021). Student-Authored Scientist Spotlights: Investigating the Impacts of Engaging Undergraduates as Developers of Inclusive Curriculum through a Service-Learning Course. CBE—Life Sciences Education, 20(4), ar55. 10.1187/cbe.21-03-0060

Ariely, M., Livnat, Z., & Yarden, A. (2019). Analyzing the Language of an Adapted Primary Literature Article: Towards a Disciplinary Approach of Science Teaching Using Texts. Science & Education, 28(1–2), 63–85. 10.1007/s11191-019-00033-5

Asterhan, C. S. C., & Resnick, M. S. (2020). Refutation texts and argumentation for conceptual change: A winning or a redundant combination? Learning and Instruction, 65, 101265. 10.1016/j.learninstruc.2019.101265

Brandt, S., Cotner, S., Koth, Z., & McGaugh, S. (2020). Scientist Spotlights: Online assignments to promote inclusion in Ecology and Evolution. Ecology and Evolution, 10(22), 12450– 12456. 10.1002/ece3.6849

Braun, V., & Clarke, V. (2006). Using thematic analysis in psychology. Qualitative Research in Psychology, 3(2), 77–101. 10.1191/1478088706qp063oa

Brown, N. J. S., Furtak, E. M., Timms, M., Nagashima, S. O., & Wilson, M. (2010). The Evidence-Based Reasoning Framework: Assessing Scientific Reasoning. Educational Assessment, 15(3–4), 123–141. 10.1080/10627197.2010.530551

Brownell, S. E., & Tanner, K. D. (2012). Barriers to Faculty Pedagogical Change: Lack of Training, Time, Incentives, and…Tensions with Professional Identity? CBE—Life Sciences Education, 11(4), 339–346. 10.1187/cbe.12-09-0163

Carter, A. W. (2018). Short heatwaves during fluctuating incubation regimes produce females under temperature-dependent sex determination with implications for sex ratios in nature. Scientific Reports, 8(1), 3.

Chi, M. T. H., & Wylie, R. (2014). The ICAP Framework: Linking Cognitive Engagement to Active Learning Outcomes. Educational Psychologist, 49(4), 219–243. 10.1080/00461520.2014.965823

Christopher Strenta, A., Elliott, R., Adair, R., Matier, M., & Scott, J. (1994). Choosing and leaving science in highly selective institutions. Research in Higher Education, 35(5), 513–547. 10.1007/BF02497086

Coil, D., Wenderoth, M. P., Cunningham, M., & Dirks, C. (2010). Teaching the Process of Science: Faculty Perceptions and an Effective Methodology. CBE—Life Sciences Education, 9(4), 524–535. 10.1187/cbe.10-01-0005

Coley, J. D., & Tanner, K. (2015). Relations between Intuitive Biological Thinking and Biological Misconceptions in Biology Majors and Nonmajors. CBE—Life Sciences Education, 14(1), ar8. 10.1187/cbe.14-06-0094

Coley, J. D., & Tanner, K. D. (2012). Common Origins of Diverse Misconceptions: Cognitive Principles and the Development of Biology Thinking. CBE—Life Sciences Education, 11(3), 209–215. 10.1187/cbe.12-06-0074

Core Team. (2022). *RStudio: Integrated Development Environment for R.* [Computer software]. R Foundation for Statistical Computing, Vienna, Austria. https://www.R-project.org/

Department of Agriculture and Forestry Service. (2013). Natural Inquirer. http://www.naturalinquirer.org/

Ebert-May, D., Derting, T. L., Hodder, J., Momsen, J. L., Long, T. M., & Jardeleza, S. E. (2011). What We Say Is Not What We Do: Effective Evaluation of Faculty Professional Development Programs. BioScience, 61(7), 550–558. 10.1525/bio.2011.61.7.9

Falk, H., Brill, G., & Yarden, A. (2008). Teaching a Biotechnology Curriculum Based on Adapted Primary Literature. International Journal of Science Education, 30(14), 1841– 1866. 10.1080/09500690701579553

Goudsouzian, L. K., & Hsu, J. L. (2023). Reading Primary Scientific Literature: Approaches for Teaching Students in the Undergraduate STEM Classroom. CBE—Life Sciences Education, 22(3), es3. 10.1187/cbe.22-10-0211

Gouvea, J. S., & Simon, M. R. (2018). Challenging Cognitive Construals: A Dynamic Alternative to Stable Misconceptions. CBE—Life Sciences Education, 17(2), ar34. 10.1187/cbe.17-10-0214

Gregory, T. R. (2009). Understanding Natural Selection: Essential Concepts and Common Misconceptions. Evolution: Education and Outreach, 2(2), 156–175. 10.1007/s12052-009-0128-1

Halmo, S. M., Bremers, E. K., Fuller, S., & Stanton, J. D. (2022). “Oh, that makes sense”: Social Metacognition in Small-Group Problem Solving. CBE—Life Sciences Education, 21(3), ar58. 10.1187/cbe.22-01-0009

Hoskins, S. G., Stevens, L. M., & Nehm, R. H. (2007). Selective Use of the Primary Literature Transforms the Classroom Into a Virtual Laboratory. Genetics, 176(3), 1381–1389. 10.1534/genetics.107.071183

Janick-Buckner, D. (1997). Getting Undergraduates to Critically Read and Discuss Primary Literature: Cultivating Students’ Analytical Abilities in an Advanced Cell Biology Course. Journal of College Science Teaching, 27(1), 29–32.

Kararo, M., & McCartney, M. (2019). Annotated primary scientific literature: A pedagogical tool for undergraduate courses. PLOS Biology, 17(1), e3000103. 10.1371/journal.pbio.3000103

Koomen, M. H., Weaver, S., Blair, R. B., & Oberhauser, K. S. (2016). Disciplinary literacy in the science classroom: Using adaptive primary literature. Journal of Research in Science Teaching, 53(6), 847–894. 10.1002/tea.21317

Kozeracki, C. A., Carey, M. F., Colicelli, J., & Levis-Fitzgerald, M. (2006). An Intensive Primary-Literature–based Teaching Program Directly Benefits Undergraduate Science Majors and Facilitates Their Transition to Doctoral Programs. CBE—Life Sciences Education, 5(4), 340–347. 10.1187/cbe.06-02-0144

Krontiris-Litowitz, J. (2013). Using Primary Literature to Teach Science Literacy to Introductory Biology Students. Journal of Microbiology & Biology Education, 14(1), 66–77. 10.1128/jmbe.v14i1.538

Kuhn, D. (1993). Science as argument: Implications for teaching and learning scientific thinking. Science Education, 77(3), 319–337. 10.1002/sce.3730770306

McCartney, M., Childers, C., Baiduc, R. R., & Barnicle, K. (2018). Annotated Primary Literature: A Professional Development Opportunity in Science Communication for Graduate Students and Postdocs. Journal of Microbiology & Biology Education, 19(1), 10–1128. 10.1128/jmbe.v19i1.1439

Meaders, C. L., Lane, A. K., Morozov, A. I., Shuman, J. K., Toth, E. S., Stains, M., Stetzer, M. R., Vinson, E., Couch, B. A., & Smith, M. K. (2020). Undergraduate Student Concerns in Introductory STEM Courses: What They Are, How They Change, and What Influences Them. Journal for STEM Education Research, 3(2), 195–216. 10.1007/s41979-020-00031-1

Mervis, J. (2011). Weed-Out Courses Hamper Diversity. Science, 334(6061), 1333–1333. 10.1126/science.334.6061.1333

Ovid, D., Abrams, L., Carlson, T., Dieter, M., Flores, P., Frischer, D., Goolish, J., Bernt, M. L.-F., Lancaster, A., Lipski, C., Luna, J. V., Luong, L. M. C., Mullin, M., Newman, M. J., Quintero, C., Reis, J., Robinson, F., Ross, A. J., Simon, H., … Tanner, K. D. (2023). Scientist Spotlights in Secondary Schools: Student Shifts in Multiple Measures Related to Science Identity after Receiving Written Assignments. CBE—Life Sciences Education, 22(2), ar22. 10.1187/cbe.22-07-0149

Petersen, C. I., Baepler, P., Beitz, A., Ching, P., Gorman, K. S., Neudauer, C. L., Rozaitis, W., Walker, J. D., & Wingert, D. (2020). The Tyranny of Content: “Content Coverage” as a Barrier to Evidence-Based Teaching Approaches and Ways to Overcome It. CBE—Life Sciences Education, 19(2), ar17. 10.1187/cbe.19-04-0079

Pickett, S. B., Nielson, C., Marshall, H., Tanner, K. D., & Coley, J. D. (2022). Effects of Reading Interventions on Student Understanding of and Misconceptions about Antibiotic Resistance. Journal of Microbiology & Biology Education, 23(1), e00220–21. 10.1128/jmbe.00220-21

Richard, M., Coley, J. D., & Tanner, K. D. (2017). Investigating Undergraduate Students’ Use of Intuitive Reasoning and Evolutionary Knowledge in Explanations of Antibiotic Resistance. CBE—Life Sciences Education, 16(3), ar55. 10.1187/cbe.16-11-0317

Round, J. E., & Campbell, A. M. (2013). Figure Facts: Encouraging Undergraduates to Take a Data-Centered Approach to Reading Primary Literature. CBE—Life Sciences Education, 12(1), 39–46.

Sakaluk, S. K., Oldzej, J., Poppe, C. J., Harper, J. L., Rines, I. G., Hampton, K. J., Duffield, K. R., Hunt, J., & Sadd, B. M. (2019). Effects of inbreeding on life-history traits and sexual competency in decorated crickets. Animal Behaviour, 155, 241–248. 10.1016/j.anbehav.2019.05.027

Saldaña, J. (2013). The coding manual for qualitative researchers (2nd ed). SAGE.

Schinske, J. N., Perkins, H., Snyder, A., & Wyer, M. (2016). Scientist Spotlight Homework Assignments Shift Students’ Stereotypes of Scientists and Enhance Science Identity in a Diverse Introductory Science Class. CBE—Life Sciences Education, 15(3), ar47. 10.1187/cbe.16-01-0002

Seymour, E., & Hewitt, N. M. (1997). Talking about leaving. (Vol. 34). Westview Press.

Shadle, S. E., Marker, A., & Earl, B. (2017). Faculty drivers and barriers: Laying the groundwork for undergraduate STEM education reform in academic departments. International Journal of STEM Education, 4(1), 8. 10.1186/s40594-017-0062-7

Stern, F., Kampourakis, K., Huneault, C., Silveira, P., & Müller, A. (2018). Undergraduate Biology Students’ Teleological and Essentialist Misconceptions. Education Sciences, 8(3), 135. 10.3390/educsci8030135

Tippett, C. D. (2010). Refutation text in science education: A Review of two decades of research. International Journal of Science and Mathematics Education, 8(6), 951–970. 10.1007/s10763-010-9203-x

Tobin, K. B., Calhoun, A. C., Hallahan, M. F., Martinez, A., & Sadd, B. M. (2019). Infection Outcomes are Robust to Thermal Variability in a Bumble Bee Host–Parasite System. Integrative and Comparative Biology, 59(4), 1103–1113. 10.1093/icb/icz031

Van Lacum, E., Ossevoort, M., Buikema, H., & Goedhart, M. (2012). First Experiences with Reading Primary Literature by Undergraduate Life Science Students. International Journal of Science Education, 34(12), 1795–1821. 10.1080/09500693.2011.582654

Watkins, J., & Mazur, E. (2013). Retaining Students in Science, Technology, Engineering, and Mathematics (STEM) Majors. Journal of College Science Teaching, 42(5), 36–41.

Yarden, A., Brill, G., & Falk, H. (2001). Primary literature as a basis for a high-school biology curriculum. Journal of Biological Education, 35(4), 190–195. 10.1080/00219266.2001.9655776

Yarden, A., Norris, S. P., & Phillips, L. M. (2015). Adapted Primary Literature: The Use of Authentic Scientific Texts in Secondary Schools (Vol. 22). Springer Netherlands. 10.1007/978-94-017-9759-7

Yonas, A., Sleeth, M., & Cotner, S. (2020). In a “Scientist Spotlight” Intervention, Diverse Student Identities Matter. Journal of Microbiology & Biology Education, 21(1), 25. 10.1128/jmbe.v21i1.2013

